# Differential glucose uptake by endothelial cells in response to in-vitro Herpes Virus 8 infection, when challenged with serums from diabetes mellitus type 2 patients

**DOI:** 10.64898/2026.03.02.708961

**Authors:** Raffaello Pompei

## Abstract

**T**his work explores the relationship between glucose uptake by human endothelial cells and infection by Human Herpesvirus 8 (HHV8). The results indicate that HHV8-infected endothelial cells uptake significantly more glucose than uninfected cells. In addition, when the endothelial cells are treated with diabetes type 2 serums (DM2), the uptake of glucose is even greater in HHV8 infected cells, but it is significantly depressed in not-infected cells. The authors conclude that HHV8 infection could play a role in metabolic modifications which characterize DM2 patients.

## INTRODUCTION

The Human Herpesvirus 8 (HHV8), is believed to be the cause of some malignant disorders, namely Kaposi’s sarcoma (KS), primary effusion lymphoma and multicentric Castleman’s disease [1-4]. HHV8 has a specific tropism to B-lymphocytes and human endothelial cells [5]. After a lytic reproductive phase at the first infection, it establishes a latent infection for the entire life span of the host, with occasional reactivation of the acute infection [6, 7]. The HHV8 latency nuclear antigen (LANA) is known to be able to immortalize primary endothelial cells, enhance cell survival in critical conditions and induce the persistence of viral DNA in the cells as an episome [8-10]. HHV8-infection has been reported to induce profound changes in the behavior of human umbilical vein endothelial cells (HUVEC) [8, 11-14]. Specifically, upon HUVEC infection, about 30% of cell metabolites that were screened, showed to be modified, including those metabolites commonly found in many cancer cells during fatty acid synthesis and glycolysis. Wang et al. [13], Carroll et al. [11] and Rose et al. [15] reported a peculiar over-expression of the insulin receptor (IR) in KS tissues compared to normal skin, and proposed that such over-expression was required to sustain the post-confluent replication of HHV8-infected cells, thus causing the formation of multi-layered foci in cell cultures.

HHV8 is endemic in some African sub-Saharan and mediterranean countries, where it is often reported to be associated to patients suffering from some widely diffused chronic diseases, such as type 2 diabetes (DM2) and cardiovascular diseases [16-22]. Previously, we have shown that HHV8-infection enhances both insulin-uptake and glucose-consumption by HHV8 infected cells [23,24]. Also, Piras et al. investigated the prevalence of HHV8 DNA and the presence of specific antiviral antibodies in DM2 patients and in control subjects [25]. They reported the presence of anti-lytic and anti-latent antibodies in HHV8-positive DM2 subjects and in non-DM2 controls. A significant prevalence of HHV8 DNA, but not any DNA prevalence among other Herpesviruses, was found in DM2 patients. These findings provide an additional support to the hypothesis that HHV8 may be a possible risk factor for diabetes [25-30].

The present study is aimed to analyze the effect of DM2 serums, either positive or negative for HHV8 infection, on metabolic modifications occurring within the cells, focusing our attention on how the glucose uptake of HUVEC cells could be affected by viral infection in HUVEC cells either in the presence or absence of serums from HHV8 positive or negative DM2 patients. Our results provide an additional proof that HHV8-infection, and the consequent immune-response, may be in some way involved in the glucose metabolic modifications commonly found in DM2 patients.

## MATERIALS AND METHODS

HUVEC cells were infected by HHV8 as described by Ingianni et al. [24]. Briefly the cells were infected by the virus in 35 mm plates containing 2x100.000 cells for 1 h at 37°C. After 10 days TPA 20 ng/ml was added into the cell culture medium and left for 3 days in order to activate the viral infection. In some plates serums from diabetic patients were added at day +4 after TPA addiction at a final concentration of 5%. Controls with non-diabetic either HHV8 positive or negative persons were used. The cells were incubated for 1 h at 37°C, washed and then a glucose-free medium was added for 60 min at 37°C. After 1 h human insulin diluted 1/1000 was added to the plates. Finally, 0.7 mCi/ml of H^3^-radioactive 2-deoxy-D-glucose (2-DD-glucose) was added and left on cells for 15 min. Then the plates were carefully washed, the cells were lysed and read in a H^3^-scintillator.

## RESULTS

### A Serums from DM2 patients induce a strong decrease of 2-DD-glucose uptake in uninfected HUVEC cells

-The values of radioactivity found in the control sample C2 were considered as the 100% control uptake of the radioactive 2-DD-glucose (Table 1). The value found in the control C2 was almost the same as in C1. When the cells were incubated with serums from diabetic patients the results were significantly different. In the cells treated with HHV8 positive serums there was a substantial fall of 2-DD-glucose uptake, that was less than 50% in the cells treated with DM2 serums positive for HHV8 in the lytic phase (D1, p<0,01), about 16% when the serum positive for HHV8 in the latent phase was used (D2, p<0,05) and about 28% when serums from HHV8 negative diabetic patients were used (D3, p<0,05).

**Table 1.** Inhibition of 2-deoxy-D-glucose uptake in HHV8 infected or not-infected HUVEC cells incubated with serums from diabetic patients.

| **Sample** | **CPM** | **Mean** | **%** | **Difference %** |
| --- | --- | --- | --- | --- |
| D1 | 2427/1533/ | 1980 | 47.28 | -53 |
| D2 | 3083/3926 | 3504 | 83.68 | -16 |
| D3 | 2745/3267 | 3029 | 72.34 | -28 |
| C1 | 3921/4475 | 4198 | 100 | -- |
| C2 | 4411/3963 | 4187 | 100 | -- |
| **Sample** | **CPM** | **Mean** | **%** | **Difference %** |
| HD1 | 16924/7691 | 12307 | 293.93 | +50.25 |
| HD2 | 10917/10933 | 10925 | 260.92 | +33.37 |
| HD3 | 9425/9544 | 9484 | 226.51 | +15.78 |
| HC1 | 8219/6898 | 7558 | 180.51 | -8 |
| HC2 | 8203/8179 | 8191 | 195.02 | 100 |

**Notes:** C1 indicates control HUVEC cells added with HHV8-positive serum; C2 control cells with HHV8 negative control serum; D1 and D2 indicate HUVEC cells treated with HHV8-positive diabetic patient serums (D1 with HHV8 in the lytic phase and D2 in the latent phase); in D3 the plates were treated with HHV8-negative diabetic serums. In HC1,
*HC2, HD1, HD2, HD3 samples the treatment was the same as indicated previously except that the cells were infected by HHV8 as indicated in the materials and methods.*

### B 2-DD-glucose uptake in HHV8-infected HUVEC cells is strongly improved in the presence of DM2 serums

-When considering the HHV8 infected HUVEC cells, the results were even more relevant. The infected cells in HC1 and HC2 showed an uptake of 2-DD-glucose almost double as compared to uninfected C1 and C2 control cells. HC2 cells showed a little not-significant reduction in glucose uptake as compared to HC1 used as control, but HD1, HD2, HD3 showed a further strong increase of 2-DD-glucose uptake with amounts that raised to about 16% for the sample HD3, more than 30% for HD2 and over 50% for the infected cells in HD1. In all these last three cases the differences were highly significant (p< 0,01).

## DISCUSSION

In the last decades several studies investigated the relationship between HHV8 infection and type 2 diabetes. Some of these have concluded that diabetes can favor the HHV8 infection by reducing the immune response, others have found a cause-effect correlation between HHV8 infection and insulin resistance/diabetes (31-38). In this study, as expected, the incubation of HUVEC cells with serums from diabetic patients induced, a significant decrease of 2-DD-glucose uptake by HHV8 not infected cells. The reduction in glucose uptake was particularly relevant when the cells were treated with serums obtained from HHV8-positive diabetic patients during the lytic phase of infection; lesser but still significant differences were shown also in the cells treated with serums obtained during the latent phase of infection or cells treated with serums of diabetic patients who were HHV8-negative. The results were even more interesting when a similar experiment was performed while utilizing HHV8-infected HUVEC cells. In all cases there was a substantial increase of 2-DD-glucose uptake in the samples which were either left untreated, or treated with serums from DM2 patients. It is noteworthy that in this case, the higher values in 2-DD-glucose uptake were scored by the infected cells which were treated with serums collected from HHV8-positive DM2 patients, during either the lytic or the latent phase of the viral infection. The present findings indicate that anti-HHV8 serums are able to further enhance the already increased uptake of glucose previously reported by Ingianni et al. [24] and Rose et al. [15] in HHV8-infected primary endothelial cells. What is known so far is that HHV8-infected cells express a higher number of IRs on the cell membrane [15, 24]; this would justify the enhanced glucose uptake by the infected cells. However, the mechanism by which HHV8 serums further increases both insulin and glucose-uptake in infected cells, is yet to be understood. The serums used in this study were polyvalent, which supposedly may contain antibodies against both lytic and latent viral proteins. Surprisingly, the results indicate*d* that glucose-uptake is further enhanced in the HHV8-infected cells when treated with serums which are positive for HHV8.

These findings led us to speculate on a possible translation of this phenomenon *in vivo*. HHV8 is considered an oncogenic virus, that can lead to the malignant angiosarcoma KS, which is particularly aggressive in immunosuppressed subjects. During the latent infection, the HHV8 is suppressed by an efficient immune system; however, the efficiency of the immune system may decrease (i.e., due to aging or other factors), and HHV8 may reactivate into the lytic phase time after time. HHV8-infected cells, as well as cancer cells, need more energy for metabolism and progression. In this context, specific anti-HHV8 serums provide additional support to contrast the persistence and spread of the virus in the host, but at the same time depress the glucose uptake in the patient’s not infected normal cells leading to hyper-glycemia and eventually to DM2. In summary, we hypothesize that anti-HHV8 serum antibodies binding to surface-exposed viral antigens, are cause of a consequent enhancement of glucose-uptake in infected cells and a dramatic fall of glucose uptake in uninfected cells, likely through an intensified interference on glucose transport systems.

## CONCLUSION

Some years ago B. Corkey in her keynote Banting lecture [31] stated that insulin hyper-secretion is a possible initiating event of obesity and diabetes, and that discovering the causes that induce insulin hyper-secretion could help to find a definite solution for diabetes treatment. This consideration led us to speculate that the over-expression of IR together with the enhanced cellular glucose uptake induced by HHV8-infection, as indicated here, as well as in our previous works, could represent a possible initiating event for diabetes. This is true in about 50% of DM2 cases examined in our experience and in our Region. In regards to the other 50% of DM2 cases, other metabolic factors or microorganism infections should be considered. Although further work is undergoing to define this alternative and stimulating hypothesis, these evidences open a path to an extensive and intriguing further exploration.

## Notes

### Competing Interest Statement

The authors have declared no competing interest.

